# Embracing the power of genomics to inform evolutionary significant units

**DOI:** 10.1101/2025.06.14.659582

**Authors:** Jayna C. Bergman, Rebecca S. Taylor, Micheline Manseau

## Abstract

Appropriate identification of evolutionary significant units (ESUs) is essential for effective conservation planning. Genomic data has emerged as a key tool to inform ESU decisions due to the increased information resolution, yet it remains unclear how genomic data are being used in practice to identify the number of ESUs. To address this, we conducted a systematic literature review and found that genomic data are increasingly being used to suggest numbers of ESUs globally across plant and animal taxa. However, our review revealed inconsistencies in how ESUs are defined, with many studies not providing a definition at all. We also found inconsistencies in the methods used to analyze genomic data, highlighting the need for greater standardization to ensure studies adequately address all components of an ESUs. Adaptive loci, a key advantage of genomic data, need to be interpreted with caution and simply identifying these loci may lead to inflated ESU estimates. Overall, we found that 68% of studies suggested an increase in the number of ESUs, and that the amount of gene flow detected did not appear to influence this conclusion. We outline how genomic data can be used to assess the two key components of ESUs and provide recommendations for future studies aiming to identify ESUs with genomic data.

## Introduction

Identifying intraspecific units for conservation is a critical strategy for protecting diversity within species (Funk et al., 2012; Coates et al., 2018; Hohenlohe et al., 2020; Allendorf et al., 2022; Forester et al., 2022; Turbek et al., 2023; Schmidt et al., 2024). One widely used concept is Evolutionary Significant Units (ESU), which are biologically informed and often legally recognized groups of populations below the species level. ESUs are broadly recognized as having two components that can be generalized as requiring populations to be: (1) geographically or genetically discrete, and (2) representative of an important component of a species evolutionary history (see reviews by Crandall et al., 2000; Fraser and Bernatchez, 2001; Funk et al., 2012; Turbek et al., 2023). The identification of ESUs has the potential to have direct implications for a population’s legal status and management (Coates et al., 2018); inappropriate identification of these units can waste limited conservation resources (Forester et al., 2022). Therefore, carefully considering how ESUs are identified is essential for effective conservation planning.

No single definition of an ESU is universally applied in conservation (see Table 1 for alternative definitions). Ryder (1986) first introduced the term ESU for conservation purposes, describing them as populations that “possess genetic attributes significant for the present and future generations of the species in question”. Building on this, Waples (1991) emphasized two components that required populations to be “substantially reproductively isolated” and to “represent an important component in the evolutionary legacy of the species”. Later, Moritz (1994) argued for the use of specific genetic markers, defining an ESU as “reciprocally monophyletic for mtDNA alleles and to show significant divergence of allele frequencies at nuclear loci”. Most recently, Funk et al., (2012) were the first to explicitly incorporate the use of genomic data and suggested that ESUs should be “historically isolated from each other and likely have important adaptive differences among them”.

**Table 1.** Definitions of Evolutionary Significant Units (ESUs) from previous publications. This is not an exhaustive list but rather shows the reference and the corresponding definitions that were cited most frequently by the 13030 publications included in this review.

| Reference | Definition | Number of times cited* |
| --- | --- | --- |
| **Ryder, 1986** | "a subset of the more inclusive entity species, which possess genetic attributes significant for the present and future generations of the species in question" | 144 |
| **Waples, 1991**<br>**(Endangered Species Act in the United States)** | "A population must satisfy two criteria to be considered an ESU: (1) it must be substantially reproductively isolated from other conspecific population units, and (2) it must represent an important component in the evolutionary legacy of the species." | 99 |
| **Moritz, 1994** | "ESUs should be (1) reciprocally monophyletic for mtDNA alleles and (2) show significant divergence of allele frequencies at nuclear loci." | 233 |
| **Crandall et al., 2000** | "...suggest that the terminology [ESU] is abandoned and replaced with a more holistic concept of species, consisting of populations with varying levels of gene flow evolving through drift and natural selection" | 77 |
| **Fraser and Bernatchez, 2001** | "A lineage demonstrating highly restricted gene flow from other such lineages within the higher organizational level (lineage) of the species" | 99 |
| **Funk et al., 2012** | "Major, intraspecific units that (1) have been historically isolated from each other and (2) that likely have important adaptive differences among them" | 100 |
| **Committee on the Status of Endangered Wildlife in Canada (Designatable Units: COSEWIC, 2023)** | "A unit below the level of a taxonomic species if it has attributes that make it both (1) discrete, and (2) evolutionary significant, where discrete means that there is currently very little transmission of heritable information from other such units and evolutionarily significant means that the unit harbors heritable adaptive traits or an evolutionary history not found elsewhere in Canada." | 66 |
| **Other citations** | NA | 1818 |
| **Not reported** | NA | 8484 |

\*This column sums to more than the total number of studies due to several studies citing multiple references

In addition to these biological definitions, many countries have developed legally defined conservation units that conceptually overlap with ESUs (Coates et al., 2018; Lehnert et al., 2023). These units are officially designated to implement policies for protection, management, and conservation status. As examples, the Committee on the Status of Endangered Wildlife in Canada (COSEWIC) used Designatable Units (DUs) and their own criteria to inform the designations (COSEWIC, 2023), whereas the U.S. Endangered Species Act adopts Waples’s definition (Waples 1991) for their conservation units, referred to as Distinct Population Segments (DPS; USFWS, 1996).

Although all ESU definitions have a similar goal of preserving the unique diversity within a species, there are challenges in their application. First, the definitions vary in how strongly they emphasize isolation among populations. For example, Waples (1991) stipulates populations being “substantially reproductively isolated”, whereas Funk et al. (2012) describes populations as “historically isolated”, implying that populations may no longer be isolated. Second, the subjective wording within these definitions leaves considerable room for interpretation of the degree of divergence that is sufficient for populations to be considered as distinct ESUs. Finally, the most widely cited definitions are at least a decade, and some more than two decades old. Since these definitions were proposed, the scope and resolution of the data, as well as the analyses available to assess the data, has advanced substantially. Despite these limitations, ESUs remain widely used in conservation, underscoring the need for clearer guidance on how data and analyses can be more consistently applied.

Genomic data are emerging as a key tool to inform the identification of ESUs (Funk et al., 2012; Barbosa et al., 2018; Waples and Lindley, 2018; Funk et al., 2019; Turbek et al., 2023). Traditional genetic markers, such as mitochondrial DNA (mtDNA) and microsatellites, have been commonly used (e.g., Moritz, 1994; Kanthaswamy et al., 2006; Jensen et al., 2014). However, these markers typically target only a small number of loci (e.g., <25), providing limited genetic information that largely reflects neutral evolutionary processes (Funk et al., 2019; Barbosa et al., 2020; Ghildiyal et al., 2023; Chinna et al., 2025).

In contrast, genomic data generated through high throughput sequencing can sample thousands to millions of loci across the entire genome, offering much higher resolution on both neutral and adaptive variation (Coates et al., 2018; Ghildiyal et al., 2023; see Box 1 for an overview of genomic sequencing approaches). Using genomic data to reassess population structure previously identified with traditional genetic markers has clarified the number and distribution of distinct groups for several species (e.g., *Microtus cabrerae*, Barbosa et al., 2018; *Pseudacris crucifer*, Cairns et al., 2021; *Podocnemis lewyana*, Gallego-García et al., 2021). Thus, the unparalleled volume of information provided by genomic data has the potential to significantly advance the ability to accurately identify population structure and, in turn, ESUs (Funk et al., 2012; Coates et al., 2018), making it an essential tool for ensuring correct identification of these units.

**Genomic data** are DNA sequences or molecular markers that encompass a large and representative set of regions across the genome to provide high resolution information (Schmidt et al., 2024). Genomic data can allow for the identification of the structure and/or function of genetic variation across the genome. In this literature review we define genomic data as high-throughput sequencing that captures a large portion of the genome, using a thousand or more markers.

Genomic data can be generated by the following sequencing methods which are grouped into several categories:

1. **SNP-based genotyping** – Refers to techniques that have a set of predefined and targeted probes with specific position of single nucleotide polymorphisms (SNPs). These SNPs target variants that are based on relevance to specific individuals/populations, adaptive loci, or particular traits. We have grouped SNP chips, SNP array, SNP assays and SNP panels together because these are all types of targeted SNP-based genotyping.
2. **Reduced representation sequencing (RRS)** – Is an approach to generate genome-wide sequencing data by sequencing only digested fragments of the genome. Fragments are sampled from across the entire genome, but only a fraction of the genome is sequenced. The following sequencing types only include non-targeted RRS methods:

• ***Double digest restriction-site associated DNA sequencing (ddRAD)*** – Uses two restriction enzymes to ensure more uniform fragments of the whole genome by having two cut sites (Peterson et al., 2012). We have considered **DArTseq** methods as ddRAD because they both deploy double enzyme digestion across the genome. DArTseq uses a different system for commercial services.
• ***Genotype-by-sequencing (GBS)*** – Single enzyme digestion of the genome that uses PCR for fragment size selection.
• ***Restriction-site associated DNA sequencing (RADseq)*** – uses one enzyme to perform a single digestion of the whole genome and then randomly interrupts the digested fragments.
• ***Multiplexed Inter-simple sequence repeat genotype by sequencing (MIG-seq***) – This method does not use restriction-enzyme digestion but rather PCR-based markers that amplify a large number of genome-wide regions without prior genetic information (Suyama and Matsuki, 2015).
3. **Whole genome sequencing** – sequencing the entire genome of an organism to get a comprehensive view of the genome. There are two types of whole genome sequencing that differ by the depth of the sequencing:

• ***High coverage whole genome sequencing*** – Whole genome sequencing that typically has coverage high enough to allow genotypes to be accurately called.
• ***Low coverage whole genome sequencing*** – Whole genome sequencing resulting in a small number of reads covering the genomic loci (e.g., <10X coverage). Genotype likelihood-based methods are used with low coverage data.
4. **Exome sequencing** – Exons are protein-coding regions of the genome. Exome sequencing is a targeted sequencing approach to capture variation within these protein-coding regions.
5. **Pooled sequencing (pool-seq)** – Multiple individual DNA samples are pooled together and then sequenced in a single run. Pooled sequencing typically has a high depth of coverage (e.g., >50X) and can be cost effective (Fuentes-Pardo and Ruzzante, 2021). However individual genotypes are undistinguishable due to the process of pooling the DNA (Fuentes-Pardo and Ruzzante, 2021).
6. **Transcriptomics** – Is the characterization of RNA transcripts through RNA sequencing that is typically used to understand gene expression and regulation.

### Aims of the study

We aimed to determine how genomic data are being used to identify ESUs across taxa. To achieve this, we reviewed published literature that uses genomic data to identify and suggest ESUs for a species. Specifically, we look at the definitions and criteria used to define ESUs to identify patterns in the type of data reported and the methods used to analyze the genomic data. We also attempted to uncover how the results impacted recommendations on the number of ESUs. Finally, we provide recommendations on how genomic results can be used to inform different components of an ESU definition.

### Literature Review

We searched the Web of Science to find studies that use genomic data to inform the identification of ESUs. We included studies that were published until January 30, 2024, where the title, abstract, or keywords included the search string (genom* OR (WGS OR “whole genome” OR *seq OR GBS OR “genotyping-by-sequencing” OR SNP* OR “single nucleotide polymorphism*” OR “target capture” OR “target DNA” OR ddRAD OR “reduced representation sequencing” OR “next-generation sequencing” OR NGS OR transcript* OR *omics OR “adaptive loci” OR “adaptive locus”) AND (“conservation unit*” OR “evolutionary significant unit*” OR “evolutionarily significant unit*” OR “adaptive unit*” OR “distinct population segment*” OR “designatable unit*”). We supplemented our search with articles from Scopus using the same set of search terms. Resulting studies from both databases were combined and duplicate studies were removed. The titles and abstracts of the retrieved articles were manually screened with the assistance of ASReview (ASReview LAB developers, 2023), followed by full-text review for those that met the initial criteria.

We defined genomic data as high-throughput sequencing that captures a large portion of the genome, using a thousand or more markers (Nielsen et al., 2020; Schmidt et al., 2024). Following this definition, we excluded papers that did not use genomic data in their analyses, including studies that had datasets with fewer than 1000 SNPs, or others using mtDNA, microsatellites, or cpDNA. We also excluded studies that did not make explicit suggestions on the number of ESUs (e.g., topics unrelated to our goals such as generating a reference genome, assigning a set of individuals to existing ESUs, or species delineation without a direct link to ESUs). For synthesis papers that did not include analysis of original data, we reviewed their cited references for additional important studies. We included studies on all vertebrate, invertebrate, and plant species and did not apply any geographic restrictions. In total, our search terms identified 415 articles of potential relevance (Table S1). Of these, 126 met our inclusion criteria and were evaluated in the literature review (Table S2).

From each study, we extracted information on sampling details and the genomic datasets used from both the main text and supplementary materials, as needed. If a study included separate datasets for more than one species, each species was treated independently and assigned a separate row in the data table. This resulted in 130 data entries (Table S2). We then recorded whether an ESU definition was provided.

There are two key components in widely used ESU definitions, both of which can be informed by genomic data: (1) discreteness, and (2) evolutionary significance. We outline how genomic data can be used to assess each, following Waples (1991) ESU definition, but adapted specifically for genomic analyses.

#### Discreteness

We consider a population (or group of populations) to be discrete if there is genomic structure identified due to differences in the frequency of alleles among genomic markers. Populations may still be considered discrete even if gene flow occurs and if they are not completely reproductively isolated, as long as they show separate structure among genomic markers.

#### Evolutionary Significance

An evolutionary significant population is defined either as one that exhibits genomic patterns as a product of past evolutionary events, such as separation in glacial refugia, or as a population with unique portion of the genomic diversity of the species that would be irreplaceable if the population was lost. Either possibility should result in a population that is on a different evolutionary trajectory compared to other populations within a species. This is similar to how Waples (1991) defined a species’ “evolutionary legacy”. But, by focusing on only genomic data, we stipulate that there must be evidence in genomic patterns, rather than simply genetic inference evident as ecological or phenotypic characteristics.

While our focus is on genomic data, legal ESU designations typically consider multiple lines of evidence, including ecological data, phenotypic data, and life history traits (e.g., Waples, 1991; Waples and Lindley, 2018; Lehnert et al., 2023). By identifying how genomic data can be used, and categorizing analyses that target the separate elements of ESU definitions (see Table 2), we want to improve how they are applied in conservation. This should also help policy makers understand how variation among ESUs is affected by the genomic methods used to define them.

**Table 2.**
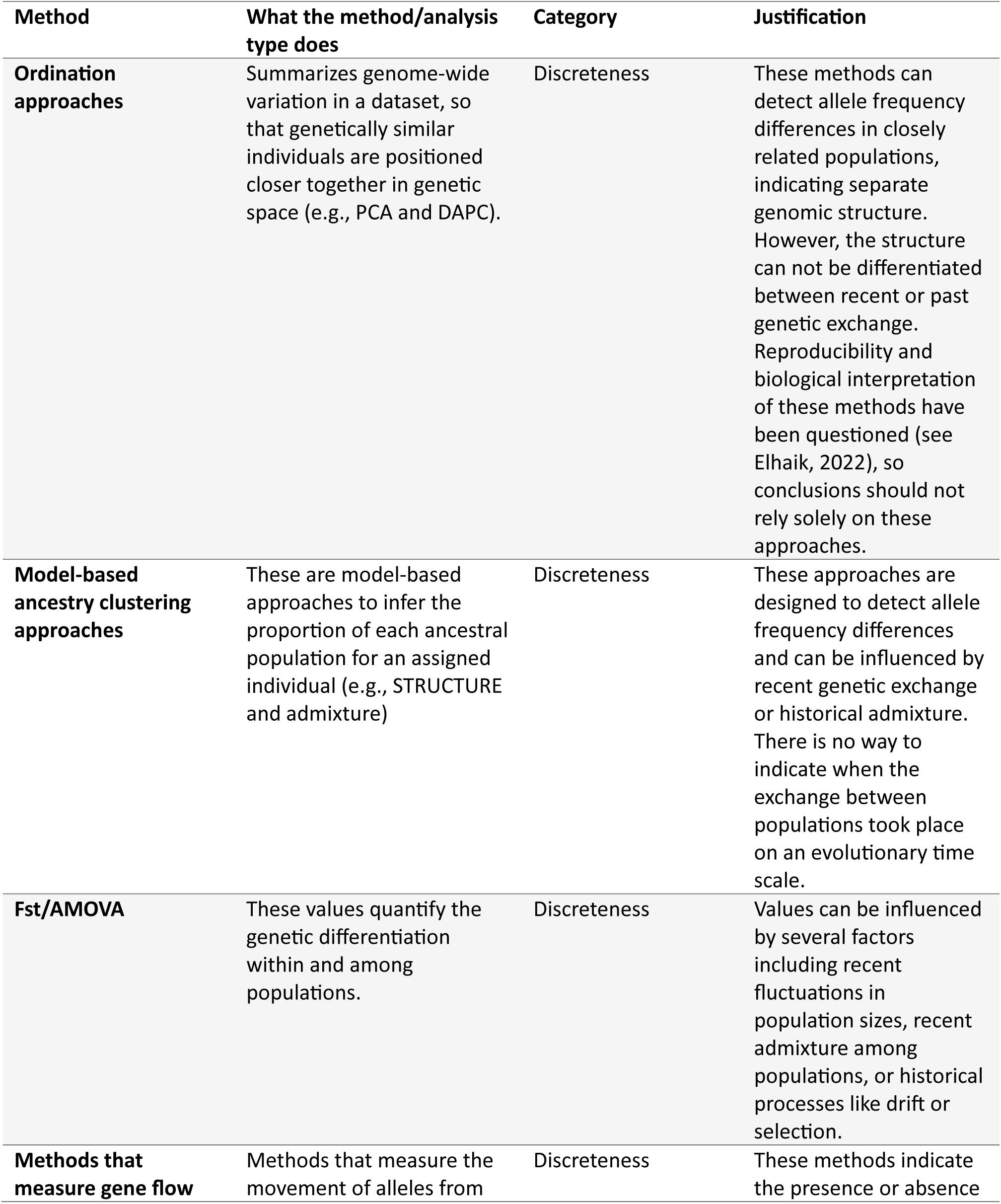
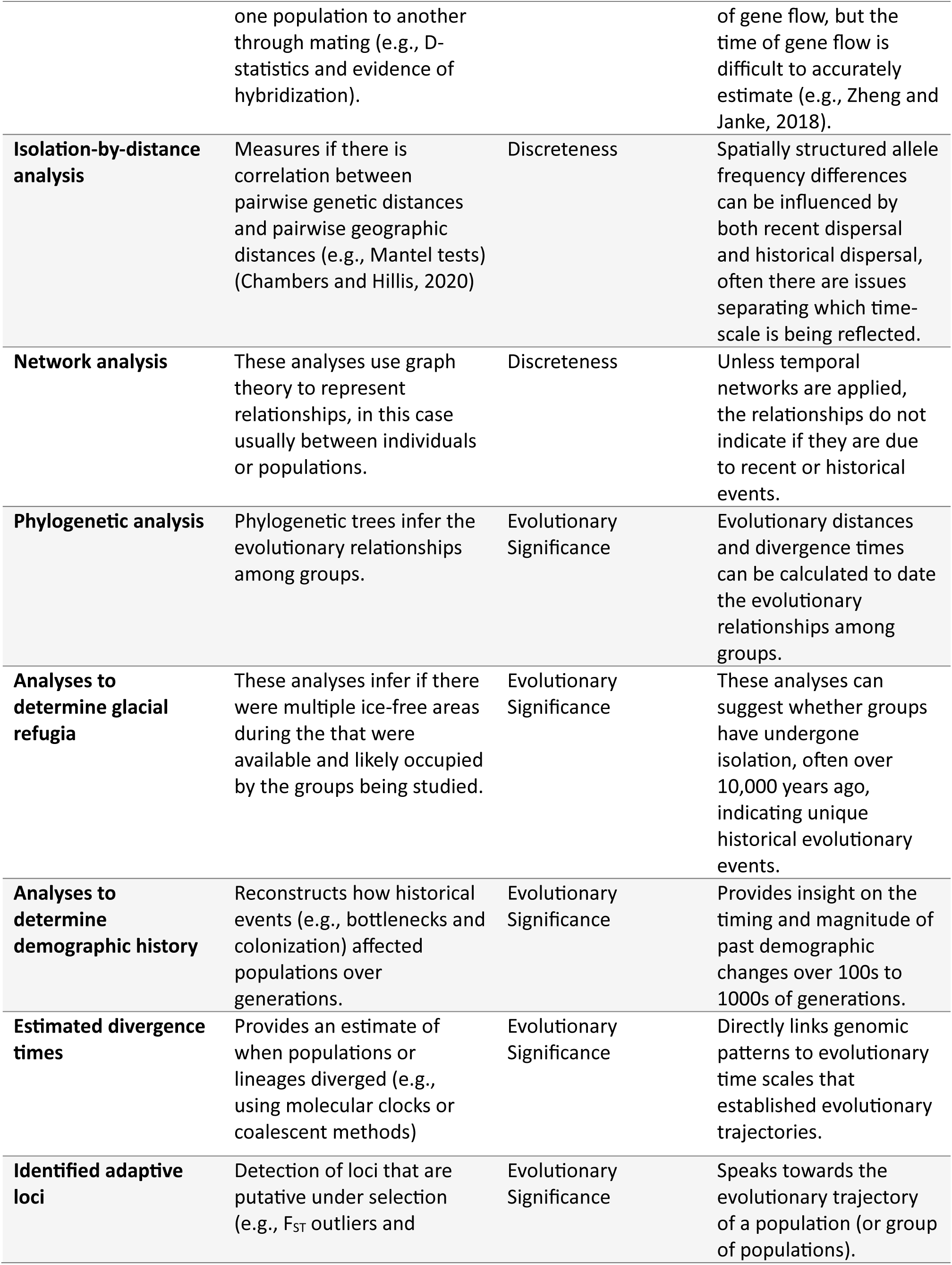
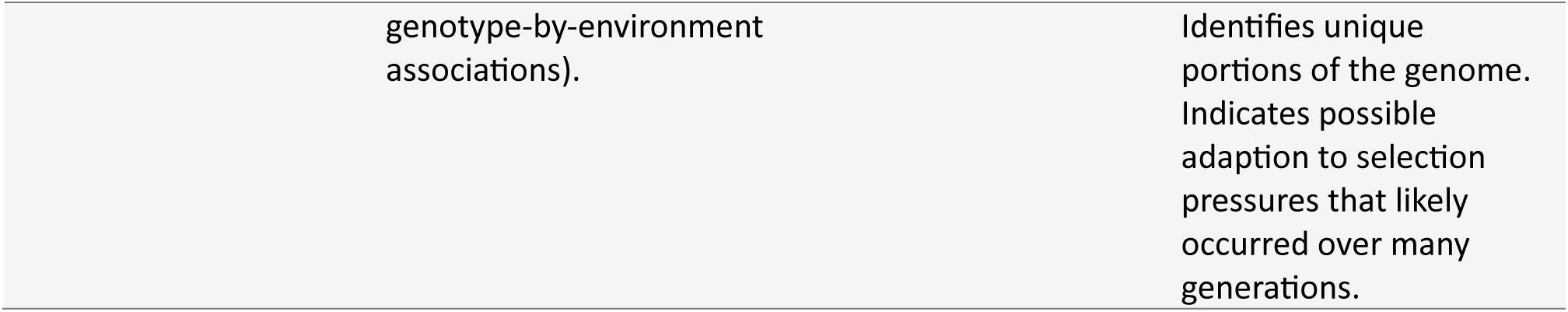
Methods used to analyze genomic data in the 130 studies included in this literature review, with justification for the categorization of each method as assessing either discreteness or evolutionary significance.

| Method | What the method/analysis type does | Category | Justification |
| --- | --- | --- | --- |
| **Ordination approaches** | Summarizes genome-wide variation in a dataset, so that genetically similar individuals are positioned closer together in genetic space (e.g., PCA and DAPC). | Discreteness | These methods can detect allele frequency differences in closely related populations, indicating separate genomic structure. However, the structure can not be differentiated between recent or past genetic exchange. Reproducibility and biological interpretation of these methods have been questioned (see Elhaik, 2022), so conclusions should not rely solely on these approaches. |
| **Model-based ancestry clustering approaches** | These are model-based approaches to infer the proportion of each ancestral population for an assigned individual (e.g., STRUCTURE and admixture) | Discreteness | These approaches are designed to detect allele frequency differences and can be influenced by recent genetic exchange or historical admixture. There is no way to indicate when the exchange between populations took place on an evolutionary time scale. |
| **Fst/AMOVA** | These values quantify the genetic differentiation within and among populations. | Discreteness | Values can be influenced by several factors including recent fluctuations in population sizes, recent admixture among populations, or historical processes like drift or selection. |
| **Methods that measure gene flow** | Methods that measure the movement of alleles from | Discreteness | These methods indicate the presence or absence |
|  | one population to another through mating (e.g., D-statistics and evidence of hybridization). |  | of gene flow, but the time of gene flow is difficult to accurately estimate (e.g., Zheng and Janke, 2018). |
| **Isolation-by-distance analysis** | Measures if there is correlation between pairwise genetic distances and pairwise geographic distances (e.g., Mantel tests) (Chambers and Hillis, 2020) | Discreteness | Spatially structured allele frequency differences can be influenced by both recent dispersal and historical dispersal, often there are issues separating which time-scale is being reflected. |
| **Network analysis** | These analyses use graph theory to represent relationships, in this case usually between individuals or populations. | Discreteness | Unless temporal networks are applied, the relationships do not indicate if they are due to recent or historical events. |
| **Phylogenetic analysis** | Phylogenetic trees infer the evolutionary relationships among groups. | Evolutionary Significance | Evolutionary distances and divergence times can be calculated to date the evolutionary relationships among groups. |
| **Analyses to determine glacial refugia** | These analyses infer if there were multiple ice-free areas during the that were available and likely occupied by the groups being studied. | Evolutionary Significance | These analyses can suggest whether groups have undergone isolation, often over 10,000 years ago, indicating unique historical evolutionary events. |
| **Analyses to determine demographic history** | Reconstructs how historical events (e.g., bottlenecks and colonization) affected populations over generations. | Evolutionary Significance | Provides insight on the timing and magnitude of past demographic changes over 100s to 1000s of generations. |
| **Estimated divergence times** | Provides an estimate of when populations or lineages diverged (e.g., using molecular clocks or coalescent methods) | Evolutionary Significance | Directly links genomic patterns to evolutionary time scales that established evolutionary trajectories. |
| **Identified adaptive loci** | Detection of loci that are putative under selection (e.g., $F_{ST}$ outliers and | Evolutionary Significance | Speaks towards the evolutionary trajectory of a population (or group of populations). |

We recorded the methods (or groups of methods) used to analyze the genomic data as either included or not, with additional columns noting the specific programs used (Table S2). Following the definitions above, all analytical methods were classified into the two broad categories as methods assessing “discreteness” and those assessing “evolutionary significance”. Methods classified under “discreteness” include ordination, model-based ancestry clustering, F_st_/AMOVA, methods that measure gene flow, isolation-by-distance analyses, and network analyses. Methods under “evolutionary significance” include phylogenetic analyses, analyses to determine demographic history, analyses to determine separate glacial refugia, estimating divergence times, and identifying adaptive loci. Justifications for why each method is in the respective category can be found in Table 2.

We assessed adaptive loci and gene flow in greater detail because both are commonly used when identifying ESUs and genomic datasets allow these methods to be applied with higher accuracy. Identifying adaptive loci, in particular, is a key advantage of using genomic data, so we were interested in assessing how these analyses were used across studies. We examined how putatively adaptive loci were identified by recording the type of analysis used to detect them and then tabulating how many studies based an ESUs evolutionary significance solely on adaptive loci identification. We also record whether the adaptive loci were simply identified as being present or if they were used as an additional dataset to further assess population structure.

Several ESU definitions incorporate gene flow and/or reproductive isolation as criteria (e.g., Waples, 1991; Fraser and Bernatchez, 2001). We therefore explored how ESUs were identified in the presence of gene flow information by noting the type of analysis used to determine it. Based on the language used by the authors in the study’s conclusions, we grouped the amount of gene flow into three categories: high, limited, or none. We then matched these categories to whether a study concluded an increase, decrease, or no change in the recommended number of ESUs (see below). This allowed us to assess whether certain ESU conclusions were consistently associated with gene flow amount.

Finally, we assessed the impact of genomic data on the number of suggested ESUs. For each study we recorded the authors’ conclusions regarding the number of ESUs and whether this represented an increase, decrease, or no change relative to the number of previously identified ESUs. If no ESUs had been previously identified for a given species, we considered it to have one ESU. We relied on the authors stated conclusions without further interpretation. Importantly, we recognize that these are theoretical suggestions rather than legal designations, but our aim was to evaluate whether the use of genomic data has led researchers to propose more ESUs, not whether these ESUs were then implemented as legal policy. Additional details are provided in the supplementary material on how data for each column was extracted from the studies (Table S3). All plots were generated in R (R Core Team, 2024) using *ggplot2* (Wickham, 2016) and *patchwork* (Pedersen, 2024).

## Results and Discussion

Our review showed a steady increase in the use of genomic data to investigate ESUs across taxonomic groups worldwide. All continents, excluding Antarctica, were represented in the dataset (Fig. 1A). However, there was a geographical bias; 57 studies (44%) were of North American species, and of those studies, 41 (32%) focused on populations within the United States (Fig. S1). Comparatively, some continents were appreciably underrepresented; only 3 studies (2%) were of African species and 7 studies (5%) on species in South America (Fig. 1A). All classes of animals were represented (Fig. 1B), however, with fish the most common (n=37, 28%) and amphibians the least (n=7, 5%). Plants were well represented in the collected studies (n=22, 17%), but a geographic bias was identified with 15 of these studies (68% of all plant studies) conducted on Asian species (Table S2). Additionally, we identified a rapid increase in the use of genomic data to identify ESUs starting in 2020 (Fig. 1C). These spatial and temporal patterns highlight the importance of understanding how these data are produced, analyzed and interpreted when applying ESUs globally.

**Figure 1.**
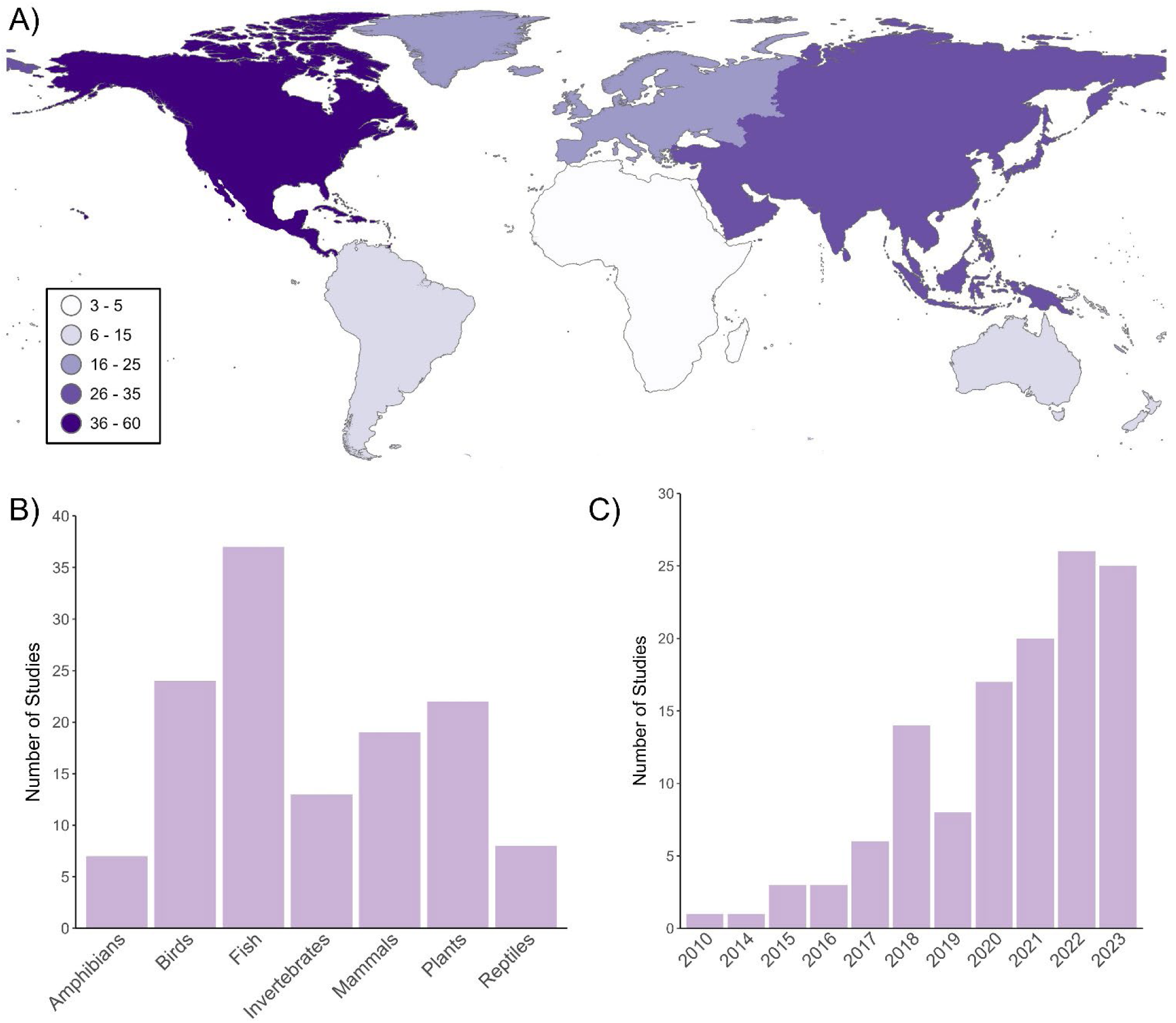
(A) Continents that are represented by the 130 studies included in the review, with darker shades indicating a higher number of studies. (B) Taxonomic groups represented in the literature review, and (C) the years the studies were published in. Our literature search ended in January 2024, and six studies from that year are included in all analyses and conclusions. However, the year 2024 is excluded from panel C because it does not represent a full year of publications.

### Definitions and criteria used to define ESUs

One of the main challenges we found was the inconsistency in how ESUs were defined across studies. Only 46 studies (35%) provided a written definition of an ESU with at least one supporting reference (Table 1). The most frequently cited definition was Moritz (1994) that was used in 23 studies (18%). Although, several other definitions were also commonly cited, including Ryder (1986), Funk et al. (2012), and Waples (1991; Table 1). Notably, many studies cited several of these references after providing a generalized definition of an ESU. Altogether, these results indicate a lack of precision in the application of specific criteria when determining whether populations (or groups of populations) should be considered as separate ESUs. Because published definitions differ, clearly identifying which ESU definition is being used is essential not only for transparency and effective application, but also to enable consistent and meaningful comparisons across studies for continuous improvement of methods.

### Genomic sequencing types used

We grouped genomic sequencing types into six broad categories (see Box 1; Fig. S2). Reduced representation sequencing (RRS) was the most common approach, used in 93 studies (72%) (RRS included only non-targeted approaches including restriction-site associated sequencing: RAD-seq, double digest restriction-site associated sequencing: ddRAD-seq, genotype by sequencing: GBS, and multiplexed inter-simple sequence repeats: MIG-seq; Fig. S2). The second most common sequencing types were whole genome sequencing (WGS), used in 21 studies (16%) and single nucleotide polymorphism (SNP)-based genotyping (e.g., SNP arrays, SNP assays, SNP chips, and SNP panels) that was used in 10 studies (8%). The remaining sequencing types included exon sequencing (n=1, <1%), transcriptomics (n=3, 2%), and pooled sequencing (pool-seq; n=1, <1%). Notably, most of these less commonly used methods target functional genomic regions and are less suited for analyses to identify population structure or evolutionary history. Furthermore, pool-seq is comparatively often confounded with a higher proportion of sequencing errors and individual genotypes are lost during the pooling process (Fuentes-Pardo and Ruzzante, 2017), making this sequencing type less effective for identifying unique population structure. Overall, RRS was used nearly five times more often than any other sequencing approach. Additionally, we found a rising trend in the use of WGS since 2018 (Fig. S3), which is expected to continue due to the increased accessibility and decreasing cost (Bagger et al., 2024; May et al., 2025).

### Data quality and reporting

We found a lack of consistent reporting on key details that are needed to assess the quality of genomic data. In particular, information on filtering decisions and their effects on the dataset is critical for evaluating data reliability and, in turn, the reliability of the results (O’Leary et al., 2018). For example, depth of coverage (also referred to as read depth) is influenced by sequencing effort and filtering decisions. Depth of coverage can affect the accuracy of genotype calls and likelihoods, and the amount of missing data, impacting downstream analyses, including population parameters estimates for inbreeding and effective population size (Hemstrom et al., 2024). Strikingly, only 57 studies (44%) reported either the average depth of coverage or the range of coverage values across samples (Table S2). Of the remaining studies, 44 (34%) reported the minimum coverage filter that was used to call genotypes, while 29 (22%) did not report any information to indicate coverage (Table S2).These reporting gaps hinder transparency and reproducibility in genomic studies.

We also observed inconsistencies in how geographic sampling information was reported. Some studies did not clearly indicate the number of geographic sampling locations, or the number of individuals sampled per location. These details are important as the number of sampling locations is strongly associated with the ability to detect patterns of isolation-by-distance (Chambers and Hillis, 2020), calculate reliable summary statistics (e.g., F_st_ values; Holsinger and Weir, 2009), and identify candidate loci for adaptation (Aguirre-Liguori et al., 2020). All of these can be useful to identify ESUs. Sampling information was not reported in eight of the studies included in our review (Table S2). While full sampling tables were often provided in the supplementary material, several studies did not summarize this information in the main text, reducing transparency and making it difficult to assess potential biases or error sources.

### Methods used to analyze genomic data

Among the various methods available to assess genomic data, the majority of studies applied more than one, although certain methods were more common (Fig. 2A). Two methods were used in nearly every study: ordination (e.g., PCA, DAPC; n=110, 86%) and model-based ancestry clustering (e.g., Admixture, Structure; n=108, 83%). These methods both evaluate discreteness and are a standard starting point for exploring population structure (see Lehnert et al., 2023; Turbek et al., 2023). In contrast, methods that assess evolutionary significance were used in fewer studies. For example, 73 (56%) generated phylogenies, 54 (42%) assessed demographic histories, and 34 (26%) estimated divergence times (Fig. 2A). Unlike discreteness, which was often assessed with multiple methods, evolutionary significance was usually assessed by a single method per study. We also found that the choice of method used to assess evolutionary significance varied across studies. This suggests that methods used to assess evolutionary significance remain less standardized than those used to assess discreteness.

**Figure 2.**
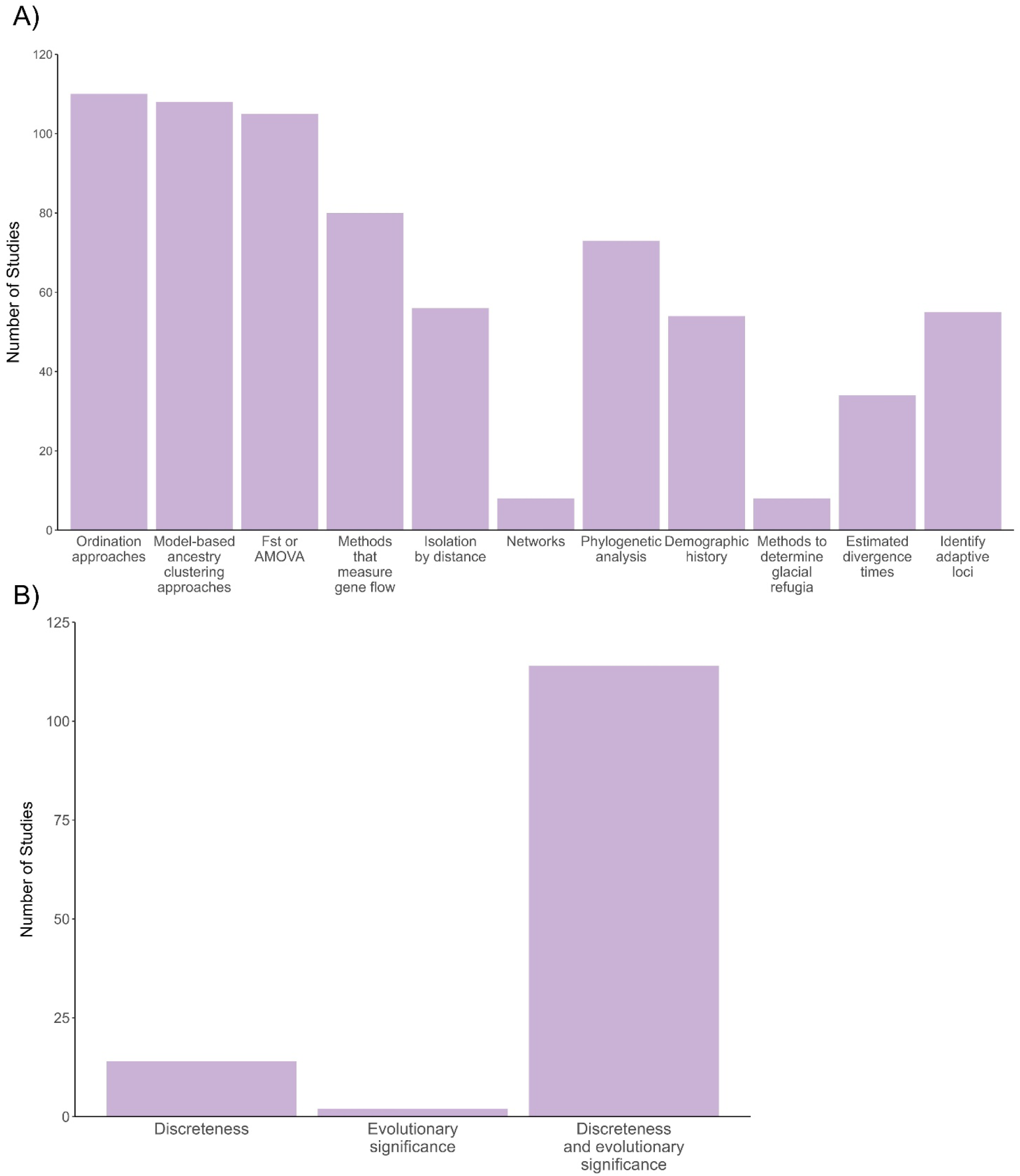
(A) The methods (or groups of methods) used to analyze genomic data across the 130 studies included in the literature review. Multiple methods were often used in each study, resulting in each bar representing the total number of times that method was used across all studies. (B) Grouping separate methods that assess the two key components of widely used evolutionary significant units (ESUs): Discreteness and Evolutionary significance. This shows the number of studies that used methods assessing only discreteness, only evolutionary significance, or using a combination of methods that assess both criteria. Methods categorized as assessing discreteness include ordination approaches, model-based ancestry clustering approaches, F_ST_ or AMOVA, methods that measure gene flow, isolation-by-distance, and network analyses. Methods categorized as assessing evolutionary significance include phylogenetic analyses, methods that assess demographic history, methods that determine glacial refugia, estimating divergence times, and identifying adaptive loci.

Despite this variability, most studies assessed both discreteness and evolutionary significance (n=114, 88%; Fig. 2B). However, 14 studies (11%) relied solely on methods that assess discreteness. This is important to note, as only assessing discreteness (e.g., ordination or model-based ancestry clustering) may produce unreliable results and inaccurately reflect a population’s evolutionary history (Lawson et al., 2018; Elhaik, 2022), potentially leading to the misidentification of ESUs. For example, inbred individuals can appear as distinct clusters on a PCA (Taylor et al., 2024) that may be misinterpreted as evidence of discrete genetic groups. In contrast, two studies (1%) relied exclusively on methods assessing evolutionary significance, and in both cases, these followed earlier studies that had already assessed discreteness. Clear and detailed examples of studies that use methods to assess both aspects of an ESU are provided in Box 2.

The following case studies are examples of thorough use of methods that address both discreteness and evolutionary significance to support the identification of ESUs using genomic data. We discuss the genomic methods applied in these studies in the context of how we have categorized methods in Table 2, which may differ from how the original authors classified their evidence. While these case studies also include non-genomic analyses, we focus our discussion on the genomic components relevant to informing ESUs.

##### Case study 1

The importance of cryptic diversity in the conservation of wide-ranging species: The red-footed tortoise (*Chelonoidis carbonarius*) in Columbia

*Gallego-Garcia et al., 2023*

The red-footed tortoise (*Chelonoidis carbonarius*) is a widespread, vulnerable species found in Central America, South America, and several Caribbean islands. Gallego-Garcia et al. (2023) sequenced 233 individuals from 25 sites across the species’ range in Colombia using triple enzyme restriction-site associated DNA sequencing (3RAD), generating a dataset of 30,327 SNPs. Although analyses were performed to determine additional smaller management units, we focus on the analyses conducted across all individuals to identify Evolutionary Significant Units (ESUs). To assess genetic discreteness, the authors used an ordination approach (Principal Component Analysis: PCA), a model-based ancestry clustering approach (STRUCTURE), and F_ST_ values. All three methods supported the presence of distinct eastern and western lineages (i.e. two ESUs). The lack of admixture in the STRUCTURE plot and high F_ST_ values (∼0.42) were used to infer no gene flow between the two groups. To assess evolutionary significance, a phylogenetic tree was generated and revealed a basal split between the lineages, supporting deep divergence and a lack of ancestral gene flow. Additionally, effective population sizes and divergence times were estimated for both lineages to further support their evolutionary significance. Finally, putatively adaptive loci were identified through genotype-environment associations (GEA) analysis using a redundancy analysis and an F_ST_ outlier analysis using PCAdapt. Overall, Gallego-Garcia et al., (2023) suggest that the eastern and western lineages should be recognized as two separate ESUs based on analyses that support both genetic discreteness and evolutionary significance, thereby increasing the number of ESUs for the species.

##### Case study 2

Selection and localised genetic structure in the threatened Manauense Harlequin frog (Bufonidae: *Atelopus manauensis*)

*Jorge et al., 2022*

The Manauense Harlequin frog (*Atelopus manauensis*) is a threaten, stream-breeding amphibian with a narrow distribution in the Amazon. Jorge et al., (2022) sequenced 94 individual males from 21 streams using a modified protocol of double digest restriction-site associated DNA sequencing (ddRAD), generating a dataset of 3,859 SNPs. To identify putatively adaptive loci, the authors used F_ST_ outlier analyses (PCAdapt and ARLEQUIN) and a genotype-environment association (GEA) approach using Latent Factor Mixed Models (LFMM). The GEA analysis detected significant associations between genetic variation and environmental variables, suggesting local adaptation. A neutral dataset of 3,121 SNPs was then created by removing the adaptive loci identified through the three analyses, as well as loci not in Hardy-Weinberg equilibrium. Discreteness was assessed using the neutral dataset through an ordination approach (Discriminant Analysis of Principal Components: DAPC), two model-based ancestry clustering approaches (sparse non-negative matrix factorization: SNMf, and STRUCTURE), and F_ST_ values. All four analyses identified six discrete genetic clusters, with high F_ST_ values (0.3-0.5) between them. To assess evolutionary significance, in addition to the initial identification of adaptive loci, the authors constructed a phylogenetic tree, and estimated divergence times, which ranged from approximately 40,000 to 2,000 years ago depending on the cluster. Finally, a stairway plot indicated population declines around 100,000 years ago, and current estimated effective population sizes per group ranged from 20 to 100 individuals. Overall, Jorge et al., (2022) suggest an increase in the number of ESUs, recognizing a total of six separate ESUs that show both genetic discreteness and evolutionary significance.

To understand whether sequencing type influenced the choice of analytical methods, we compared approaches across studies that used the three most common sequencing types: RRS, WGS, and SNP-based genotyping (Fig. 3). Across all three sequencing types, ordination and model-based ancestry clustering were the most frequently applied, appearing in at least 80% of studies for each type. F_ST_ or AMOVA methods were also commonly used in SNP-based genotyping and RRS studies (∼80%), but their use decreased to 68% in WGS studies. In contrast, we observed larger differences when comparing the methods used to assess evolutionary significance. Most notably, SNP-based studies applied these methods at much lower proportions than RRS and WGS, with the exception of identifying adaptive loci. This likely reflects the limitations of SNP-based genotyping that often leaves large portions of the genome unsampled (Dokan et al., 2021) and relies on targeted approaches that require prior knowledge of the available genetic variation (Chhina et al., 2025). These constraints can limit the ability to detect unidentified population structure or assess evolutionary significant processes.

**Figure 3.**
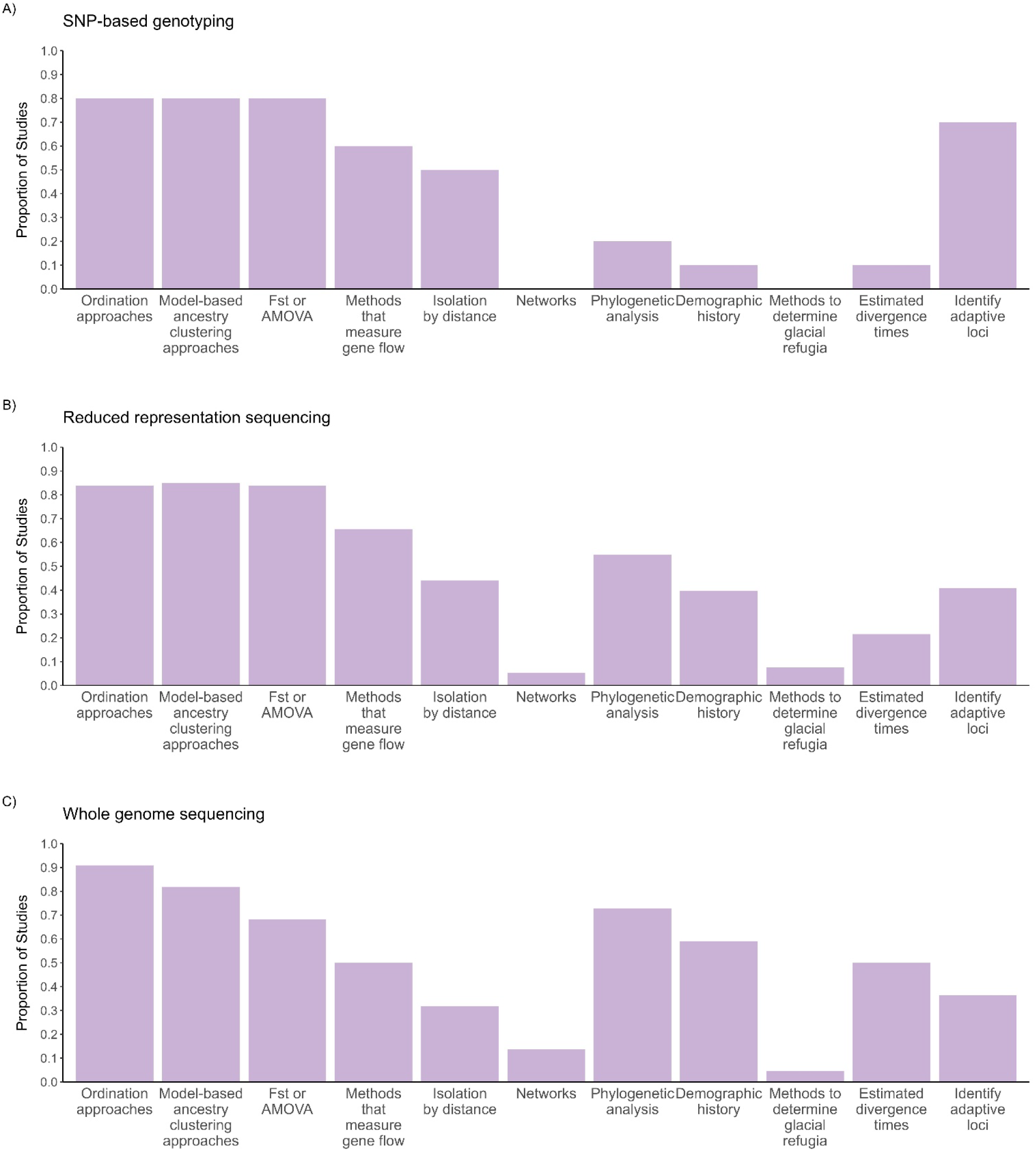
Proportion of studies using different methods to analyze genomic data from the three most commonly used sequencing types: (A) SNP-based genotyping (including SNP arrays, SNP assays, and SNP chips; used in 10 studies), (B) reduced representation sequencing (RRS; only non-targeted approaches including restriction-site associated sequencing: RAD-seq, double digest restriction-site associated sequencing: ddRAD-seq, Genotype by sequencing: GBS, and Multiplexed Inter-simple sequence repeats: MIG-seq; used in 93 studies) and (C) whole genome sequencing (WGS; used in 21 studies). Multiple methods were often used within each study.

In comparison, studies using RRS and WGS showed more similar patterns in the use of methods to assess evolutionary significance (Fig. 3). In both cases, generating a phylogeny was the most common method (RRS=55%, WGS=73%). However, approximately 20% more WGS studies used phylogenetic analyses than RRS studies, reconstructed demographic histories, and estimated divergence times. Although RRS is valid for all of these methods, the higher proportion of WGS studies may reflect a key advantage of WGS; it has the ability to provide a more comprehensive view of a species’ evolutionary history by sequencing the entire genome rather than sampling portions of it (Fuentes-Pardo and Ruzzante, 2017). Overall, these results suggest that sequencing type affects the choice of methods used to analyze genomic data, likely due to differences in data resolution.

### Adaptive genomics

A widely recognized advantage of using genomic data over mtDNA or microsatellites to identify ESUs is the improved ability to detect adaptive genome regions through the identification of adaptive (or putatively adaptive) loci (Funk et al., 2012; Barbosa et al., 2018; Waples and Lindley, 2018). In total, 55 studies (42%) identified adaptive loci. Among these studies, two approaches were used: 34 studies (62%) using F_ST_ outlier analyses and 21 studies (38%) using genotype-environment association (GEA) approaches. F_st_ outlier analyses (e.g., BayScan, Lositan) detect loci that fall outside the expected F_st_ value distribution (Funk et al., 2012) that, are often assumed to be under divergent selection and used as proxies for local adaptation (Waples and Lindley, 2018). Alternatively, GEA approaches (e.g., RDA, LFMM) identify associations between allele frequencies and environmental variables, revealing loci that are associated with specific environmental conditions (Hoban et al., 2016; Forester et al., 2018).

Of the 55 studies that identified adaptive loci, 18 (33%) based the evolutionary significance component solely on adaptive loci. Workflows to help guide ESU designation often incorporate adaptive loci identification (e.g., Funk et al., 2012; Lehnert et al., 2023; Turbek et al., 2023), which has frequently been shown to be useful to determine the evolutionary significance of populations (e.g., Barbosa et al., 2018; Waples and Lindley, 2018). However, interpreting adaptive loci requires caution (Hoban et al., 2016; Allendorf, 2017; Kardos, 2023). F_st_ outliers can result from processes other than local adaptation (Bierne et al., 2013), such as a species’ demographic history (Whitlock and Lotterhos, 2015) or genetic drift (Lotterhos and Whitlock, 2014). Similarly, loci identified through GEA methods often include false positives and may not reliably indicate local adaptation (Nadeau et al., 2016). In some cases, GEA analyses can be paired with common garden experiments or genome-wide association studies (GWAS) to verify which loci are associated with climate adaptation (e.g., Faske et al., 2021; Aitken et al., 2024), thereby strengthening the inference that these loci are truly adaptive (Hoban et al., 2016). Thus, when adaptive loci are used, studies may benefit from additional analyses that support the significance of any identified groups (e.g., phylogenetic analyses or reconstructing demographic histories).

### Gene flow

Gene flow is widely recognized as a critical evolutionary force that shapes genetic structure within and among species (Tigano and Friesen, 2016; Aitken et al., 2024; Chhina et al., 2025). It is defined as the movement of genetic information from one genetic group to another, typically through the migration of individuals (Funk et al., 2012; Tigano and Friesen, 2016). Several ESU definitions suggest that there should be substantial reproductive isolation (e.g., Waples, 1991) or little to no gene flow between groups (e.g., Fraser and Bernatchez, 2001; COSEWIC, 2023). These perspectives may reflect the concern that gene flow could erode genetic differences and reduce local adaptation (Lenormand, 2002), thereby diminishing the unique genetic diversity that defines an ESU. Alternatively, it is increasingly recognized that gene flow can occur among populations adapted to different local conditions, and that local adaptation can be maintained despite high levels of gene flow (Tigano and Friesen, 2016; Chhina et al., 2025). Other ESU perspectives emphasize the potential benefits of maintaining some level of gene flow to preserve adaptive differences (Felsenstein et al., 1976; Crandall et al., 2000). These different views highlight the absence of a unified approach and leave substantial room for subjective interpretation of how much gene flow is acceptable when recommending ESU designation.

Among the 80 studies (62%) that assessed gene flow, we found a wide range of approaches used to determine the amount of gene flow (Fig. 4A). The most common methods included interpreting admixture plots (n=13, 10%), inferring gene flow from measures of genetic differentiation (i.e., F_ST_ values; n=15, 12%), and estimating migration rates/generating migration models (n=19, 15%; Fig. 4A). While widely applied, many of these methods provide indirect or qualitative assessments rather than direct gene flow measures, which may introduce interpretive subjectivity. In contrast, more quantitative methods such as D- and F-statistics were rarely used (n=6, 5%), despite their sensitivity for detecting gene flow (Zheng and Janke, 2018). Overall, there was little consensus among studies regarding the methods used to assess gene flow.

**Figure 4.**
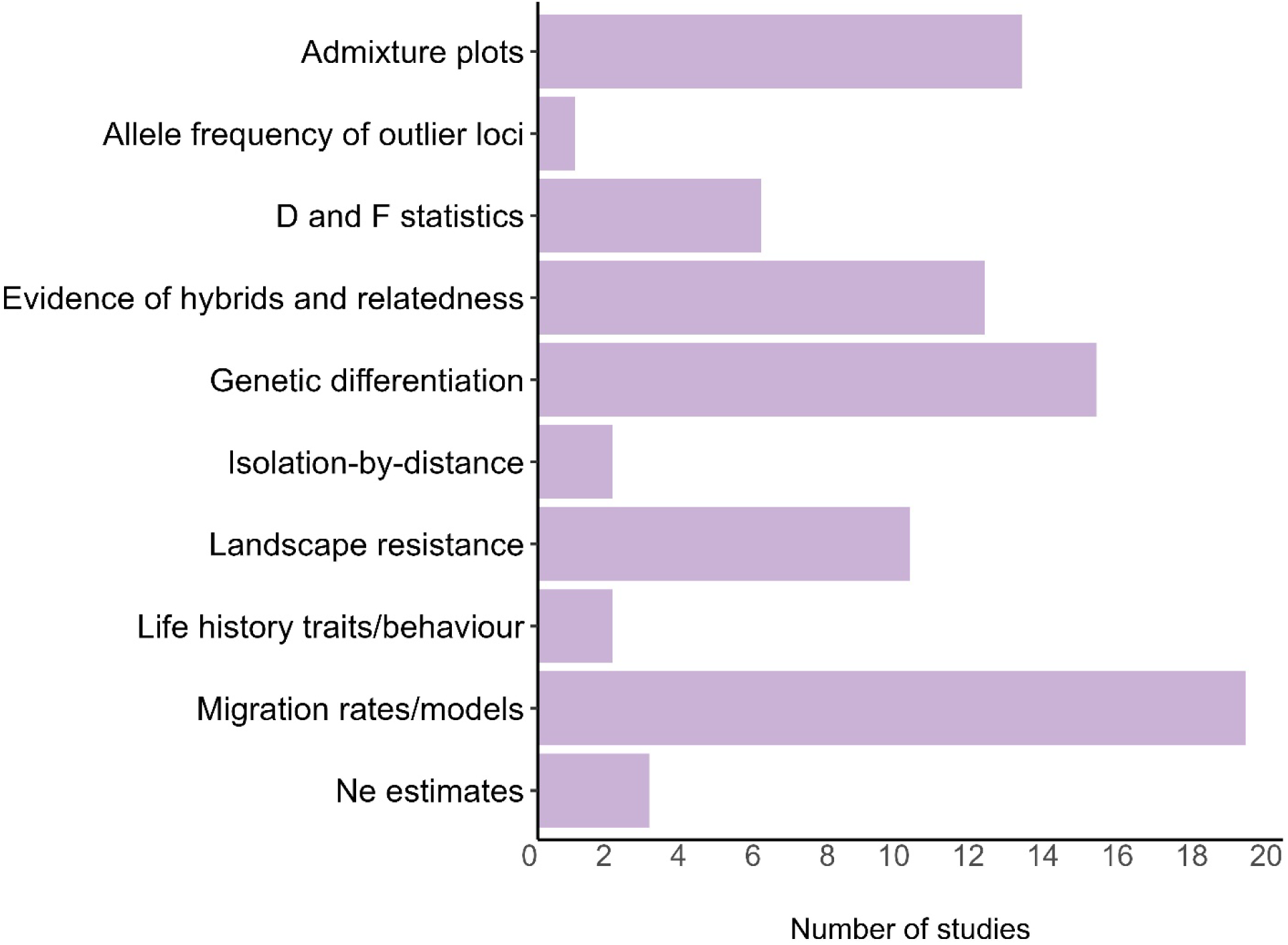
Number of studies that detected gene flow using various genomic methods. In total, 80 studies (62%) assessed gene flow among those included in the literature review.

### Changes in the number of suggested ESUs

The studies in our review more often recommended increasing the number of ESUs than maintaining the current number or decreasing them. Specifically, 89 studies (68%) suggested increasing the number of ESUs, 28 (22%) suggested no change, and only 13 (10%) proposed a decrease (Fig. S4). This trend was consistent regardless of how ESUs were defined, the sequencing type, or which downstream methods were used to analyze genomic data. These findings indicate that genomic data are generally leading researchers to identify previously undetected genetic variation within species, despite the many differences among studies. However, the proposed increases in ESUs remain theoretical suggestions and do not directly translate into legally recognized changes in ESU designations.

We examined whether the way adaptive loci were used in a study influenced the number of ESUs that were recommended. Studies using adaptive loci were categorized as either having identified adaptive loci or using them as a separate dataset in downstream analyses to assess population structure based on adaptive variation. Among the 21 studies that used GEA approaches, nine (43%) only identified the adaptive loci, whereas 12 studies (57%) incorporated them as a dataset (Fig. 5A). Studies that used adaptive loci as a dataset were more likely to suggest no change in the number of ESUs (Dataset: 4 out of 12 studies [33%]; Identified: 1 out of 9 studies [11%]; Fig. 5B). In contrast, among studies that used F_st_ outlier analyses, 23 (72%) only identified the adaptive loci, and 9 (28%) used them in downstream analyses. Although more studies only identified adaptive loci when using F_ST_ analyses, a smaller proportion of these studies suggested a decrease in the number of ESUs (Dataset: 3 out of 9 [33%]; identified: 2 out of 23 [9%]; Fig. 5B). This pattern suggests that simply identifying adaptive loci, without additionally examining their population structure, may inflate the number of ESUs identified, possibly due to the high rate of false positives associated with these methods (Bierne et al., 2013; Nadeau et al., 2016). Further analysis of adaptive loci to assess population structure may help mitigate this effect. Ultimately, while loci identified through F_st_ outliers and GEA approaches are useful proxies for local adaptation, their interpretation should be approached with caution when informing ESUs.

**Figure 5.**
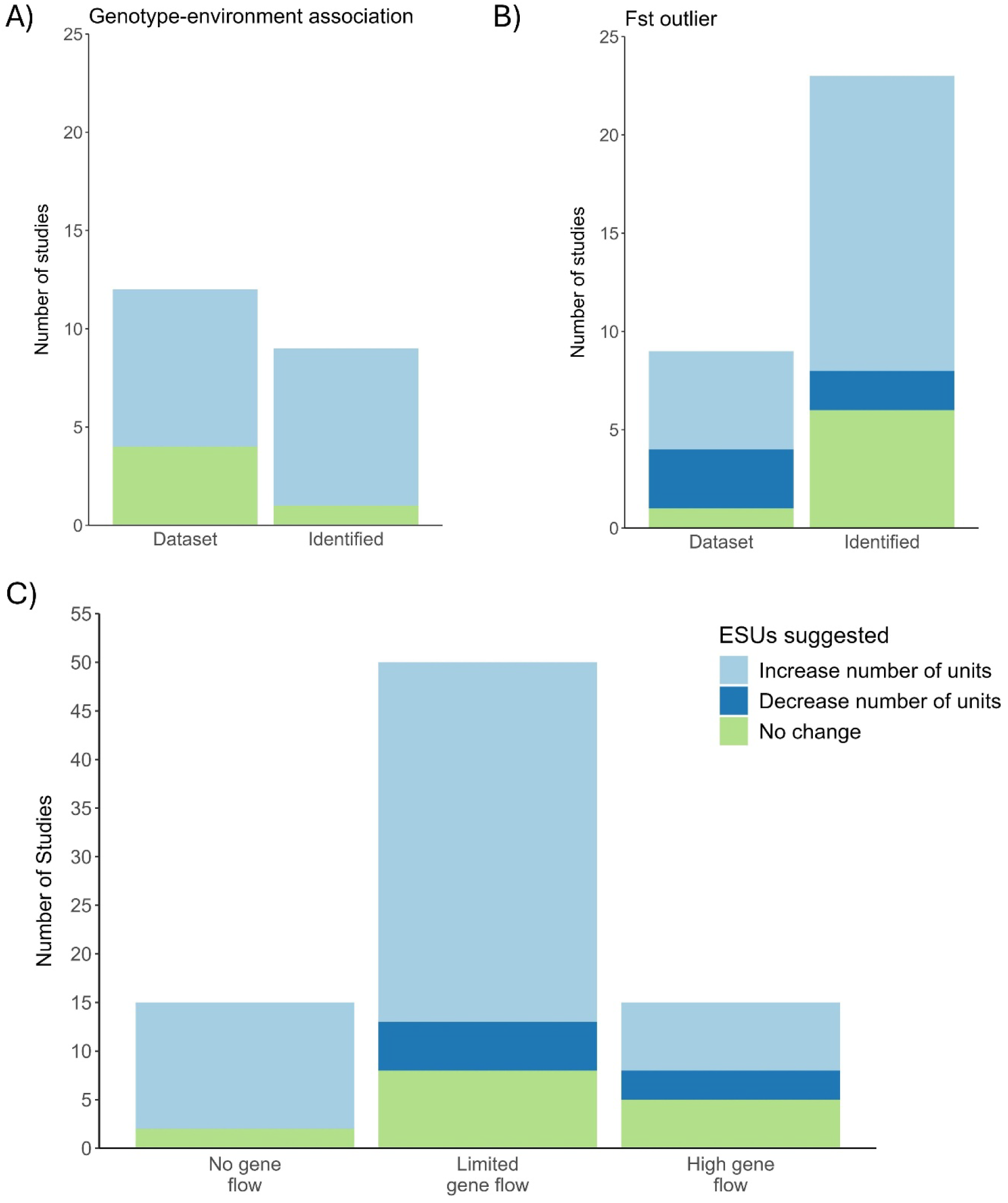
Impact of two key groups of genomic methods on the number of evolutionary significant units (ESUs) suggested by each study. (A) Studies identifying adaptive loci using genotype-environment associations (GEA) approaches, and (B) studies identifying adaptive loci using F_ST_ outlier analyses. For (A) and (B), studies are further divided based on whether adaptive loci were only identified or also used as a dataset to further assess population structure. (C) Studies that detected gene flow, grouped by the amount of gene flow reported by the authors and whether the amount of gene flow influenced the number of ESUs suggested. Colors indicate whether a study suggested an increase, decrease, or no change in the number of ESUs.

We also examined whether the amount of gene flow reported influenced the suggested number of ESUs. Based on the language used in each study’s conclusions, we categorized gene flow into three levels: high, limited, or none. We note that because the methods used varied among studies and there are no standard thresholds, these categories are arbitrary. Nevertheless, we found that they provide useful insights on how gene flow is influencing the number of ESUs being suggested. Surprisingly, the amount of gene flow reported did not appear to influence whether a study suggested increasing or decreasing the number of ESUs (Fig. 5C). Even among studies that described “high gene flow”, the largest proportion still proposed to increase the number of ESUs (Fig. 5C). This result contrasts with the current ESU definitions that emphasize the importance of limited gene flow or high amounts of reproductive isolation between groups (e.g., Waples, 1991; Fraser and Bernatchez, 2001; COSEWIC, 2023), suggesting inconsistency between the definitions and how they are being applied in research.

## Conclusion and Recommendations

The vast amount of information that genomic data provide greatly improves our ability to identify ESUs (Funk et al., 2012; Coates et al., 2018; Funk et al., 2019; Turbek et al., 2023). Genomic data facilitates a deeper understanding of the genetic differences within a species and can reliably uncover their evolutionary history, both of which are essential to accurately identify intraspecific ESUs. We conducted a comprehensive literature review to explore how genomic data have been used to suggest ESUs globally. We found that key details about the genomic data being used, such as depth of coverage, are frequently not reported. Additionally, we found that there is inconsistency in how ESUs are defined and which methods are used to analyze genomic data. Despite these gaps, most studies detected additional genetic variation leading to a recommended increase in the number of ESUs. Based on the literature review findings, we recommend future research that use genomic data to identify ESUs:

1. Define ESUs explicitly. Provide a clear definition of what is being considered an ESU in the study and link the analyses and results to the relevant components of the definition. We show how genomic data can be used to support the two key components of most ESU definitions, “discreteness” and “evolutionary significance”, to promote greater consistency in the analysis of genomic data when being used to inform ESUs.
2. Clearly report the filtering steps for the genomic data and key parameters to assess genomic data quality. For example, include the number of reads at different filtering steps (see recommendations by O’Leary et al., 2018 and Hemstrom et al., 2024), a site frequency spectrum of the final dataset as additional materials, final depth of coverage across samples, and clearly describe sampling designs and sample sizes.
3. Use multiple complementary methods to thoroughly assess population structure in ways that assess both “discreteness” and “evolutionary significance”. Avoid basing conclusions solely on measures of discreteness, particularly results from ordination or model-based ancestry clustering. These analyses can be strongly influenced by recent isolation due to habitat fragmentation, genetic drift, or inbreeding, which may result from anthropogenic causes (Schlaepfer et al., 2018).
4. Interpret adaptive loci (e.g., F_ST_ outliers) cautiously and use an additional method to confirm the evolutionary significance of groups identified through these methods. Although useful in many circumstances, F_ST_ outliers can be influenced by non-adaptive processes like sampling scheme or bioinformatic artifacts (Bierne et al., 2013; Lotterhos and Whitlock, 2014; Nadeau et al., 2016).
5. Some degree of gene flow should be allowed between ESUs, especially if this is the natural state of the populations. Gene flow among populations can maintain and even increase genetic diversity (Aitken and Whitlock, 2013; Aitken et al., 2024), and conservation management should aim to maintain this natural gene flow, even if populations are considered as separate ESUs (Chhina et al., 2025).
6. Identify ESUs in a manner that preserves a species’ evolutionary legacy. Consider using phylogenomic and demographic history reconstruction analyses to capture a species’ evolutionary history.
7. When possible, use whole genome sequencing data or dense genotype-by-sequencing SNP datasets. These dense sequencing types offer greater resolution than other sequencing types and could provide the much-needed consistency for identifying ESUs across taxa.

Despite these suggestions for improvement, significant progress has been made over the past decade to better use the power of genomics to inform ESU identification (Flanagan, et al., 2018; Funk et al., 2019; Allendorf et al., 2022). Genomic data, including WGS, are more accessible than ever before (Lou et al., 2021; Schmidt et al., 2024; May et al., 2025) and can provide valuable insights into genome-wide variation inclusive of neutral, adaptive, and deleterious variation that may be critical for current and future conservation management (Flanagan et al., 2018; Chhina et al., 2024; Schmidt et al., 2024). In addition, genomic data facilitates a wide range of analyses (Schmidt et al., 2024) that can be helpful to identify the evolutionary significant component required for populations to be considered as separate ESUs. By fully embracing genomics with more standardization and thoughtful interpretation, we can improve species conservation and ensure that decisions regarding the identification of ESUs are made accurately and informatively for evolutionary relevance.

## Acknowledgements

This research was supported by Environment and Climate Change Canada (ECCC).

## Data Availability Statement

The authors have nothing to report.

