## Supplemental Figures (1 to 4) for "Embracing the power of genomics to inform evolutionary significant units"

**Supplementary Materials**

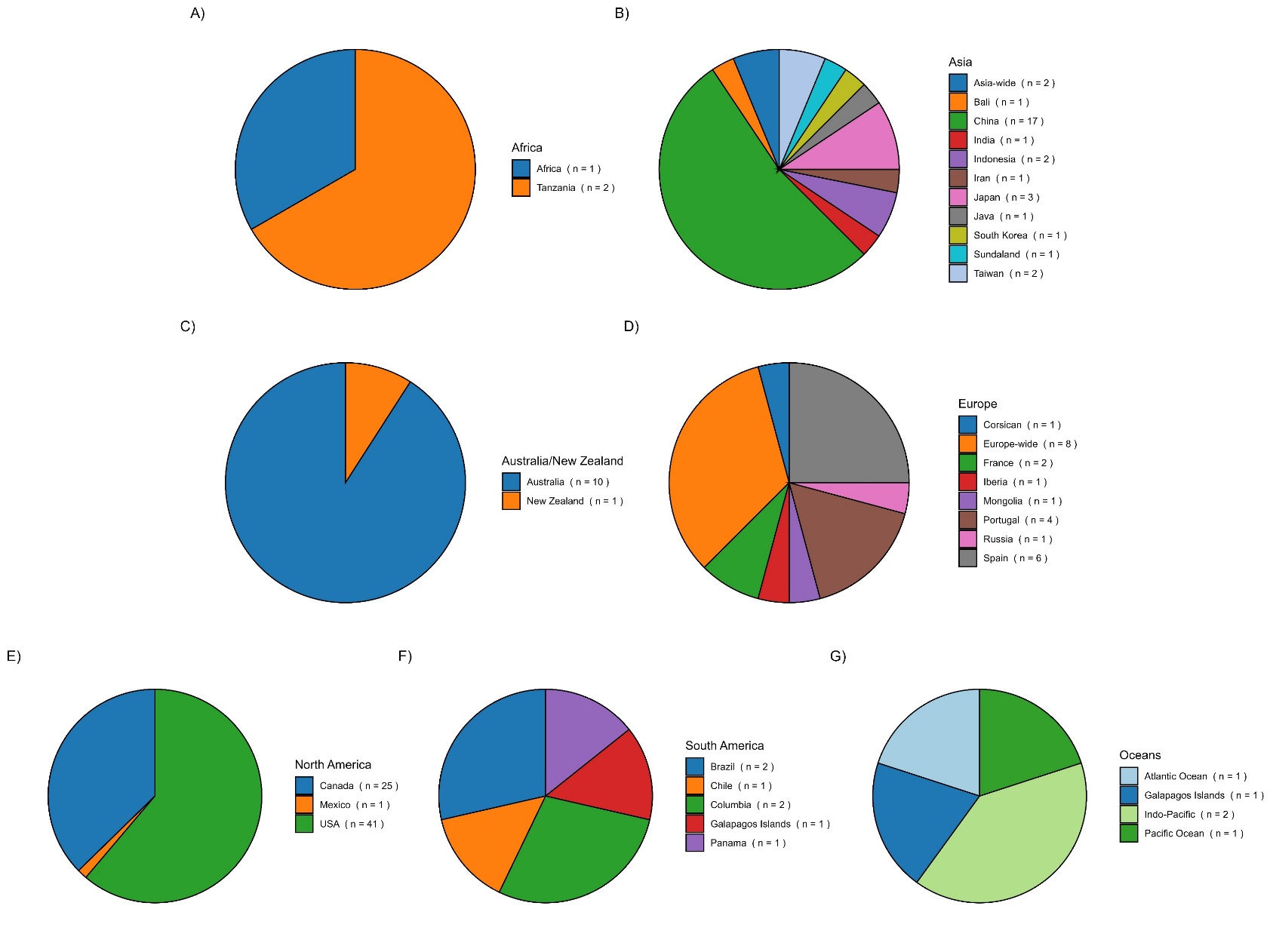

Figure S1. The number of studies that took place in each country, for the 130 studies included in this review, grouped by continent: (A) Africa, (B) Asia, (C) Australia, (D) Europe, (E) North America, (F) South America, and (G) the Oceans.

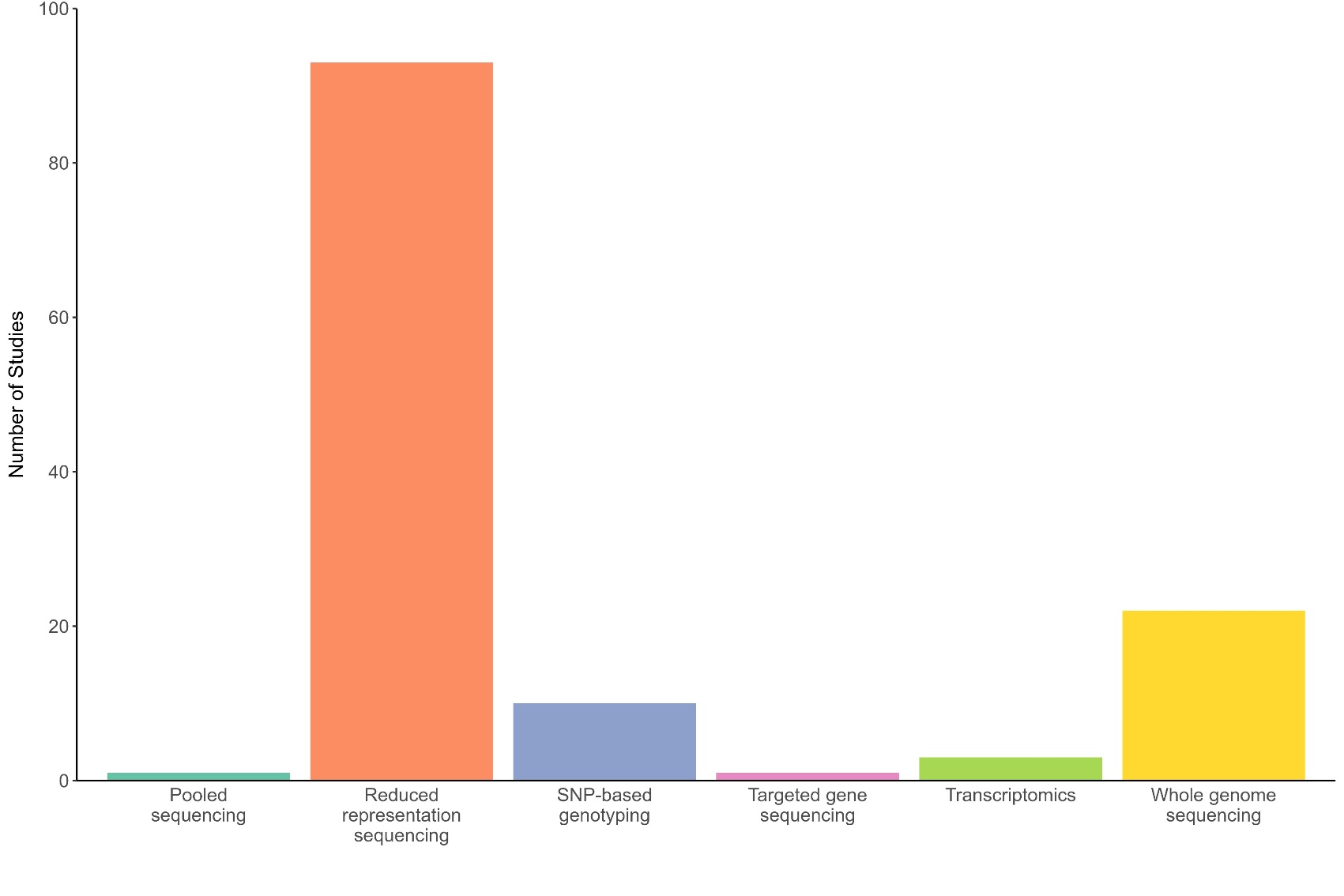

Figure S2. The number of studies that used different types of sequencing methods out of the 130 studies included in this review. Reduced representation sequencing is inclusive of only non-targeted restriction-site associated sequencing: RAD-seq, double digest restriction-site associated sequencing: ddRAD-seq, genotype by sequencing: GBS, and multiplexed inter-simple sequence repeats: MIG-seq. SNP-based genotyping is inclusive of SNP arrays, SNP assays, and SNP chips. Targeted gene sequencing includes exon sequencing.

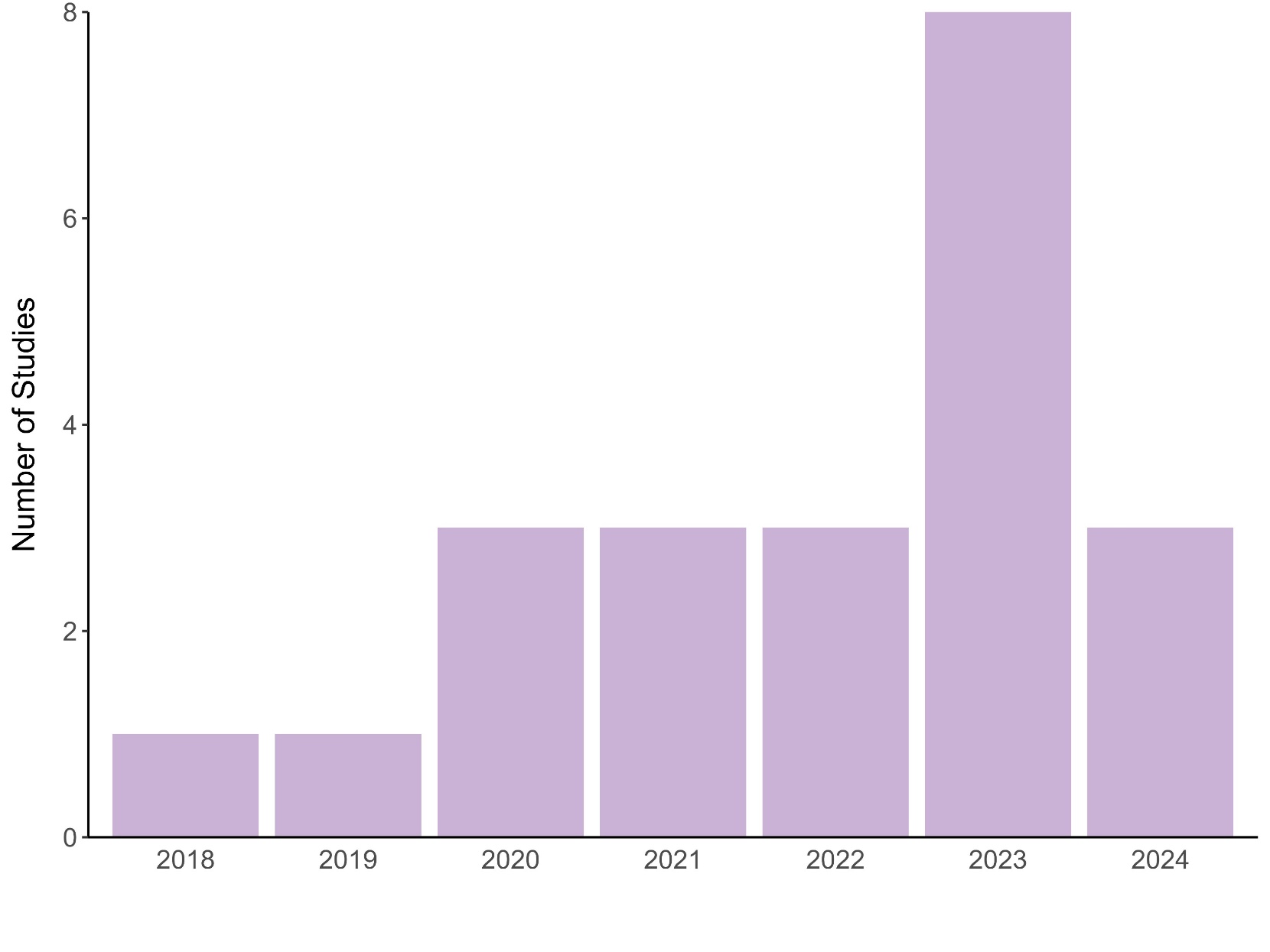

Figure S3. The number of studies using whole genome sequencing per year, based on publications included in this literature review (n=22). Studies were included up to January 2024 (total number of studies from January 2024 = 6); therefore, the count for 2024 represents only one month of data.

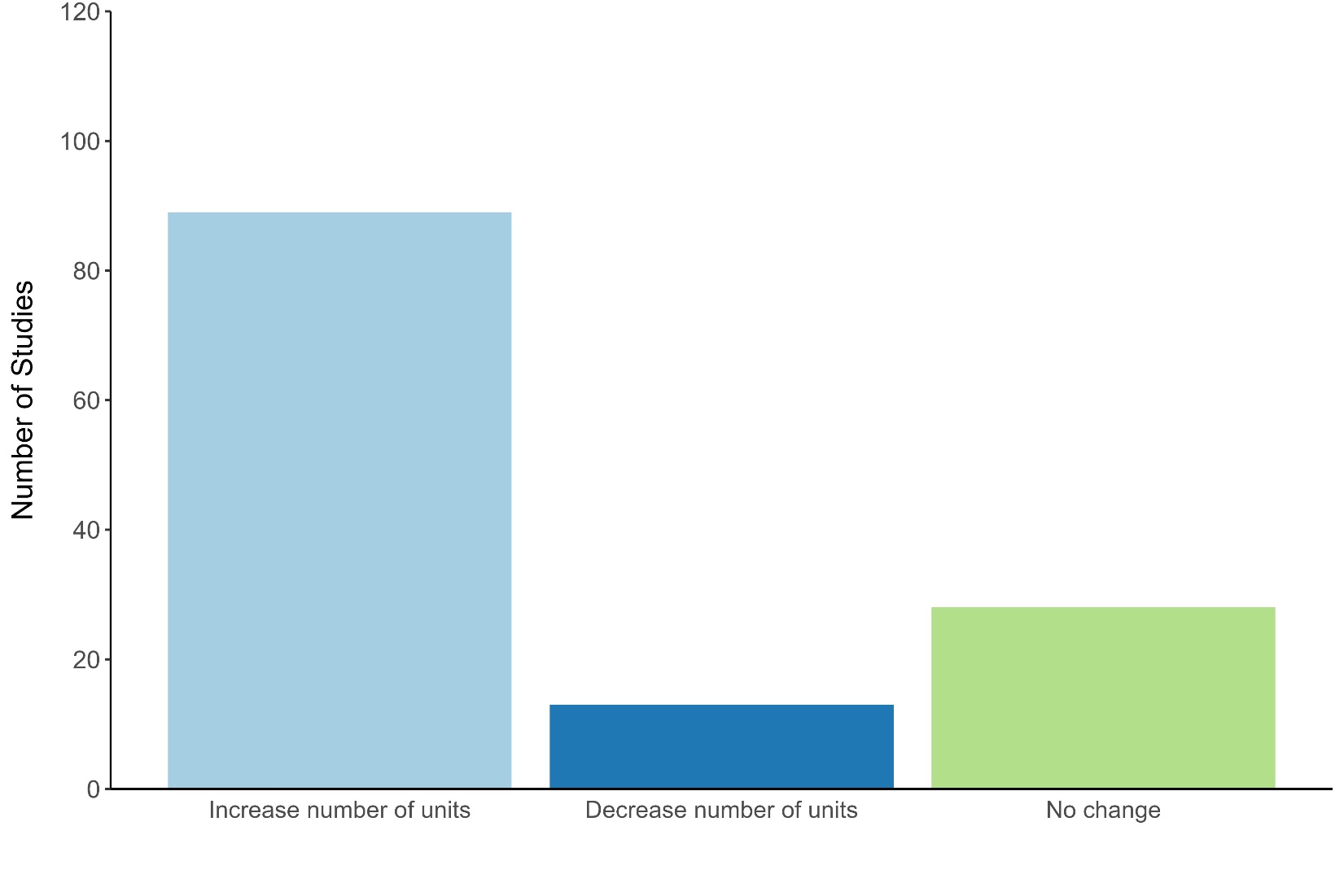

Figure S4. The overall conclusions on the number of evolutionary significant units (ESUs) that were suggested by each study as either an increase, decrease, or no change in the number of ESUs for each of the 130 studies in the literature review.
